# Spike Neural Network of Motor Cortex Model for Arm Reaching Control

**DOI:** 10.1101/2024.02.07.579412

**Authors:** Hongru Jiang, Xiangdong Bu, Xiaohong Sui, Huajin Tang, Xiaochuan Pan, Yao Chen

**Author notes:** e-mail: {, }.

## Abstract

Motor cortex modeling is crucial for understanding movement planning and execution. While interconnected recurrent neural networks have successfully described the dynamics of neural population activity, most existing methods utilize continuous signal-based neural networks, which do not reflect the biological spike neural signal. To address this limitation, we propose a recurrent spike neural network to simulate motor cortical activity during an arm-reaching task. Specifically, our model is built upon integrate-and-fire spiking neurons with conductance-based synapses. We carefully designed the interconnections of neurons with two different firing time scales - “fast” and “slow” neurons. Experimental results demonstrate the effectiveness of our method, with the model’s neuronal activity in good agreement with monkey’s motor cortex data at both single-cell and population levels. Quantitative analysis reveals a correlation coefficient 0.89 between the model’s and real data. These results suggest the possibility of multiple timescales in motor cortical control.

## I. INTRODUCTION

Motor cortex (MC) is fundamental for the generation of voluntary movements. Approaches to understand how the MC controls movement involve recording neuronal activity while animals perform tasks. Traditionally, many of these experiments focus on the representations that relate single-neuron activity to specific movement parameters, such as movement direction [1], effector position and velocity[2]. However, the representative perspective fails to explain the phenomenon where some neurons do not relate to any specific movement parameter, while others simultaneously relate to multiple movement parameters.

Recently, there has been a shift in MC studies from single-neuron representations to the dynamics that generates neural population activity. Studies have emphasized that the MC is a dynamical system, where movement preparation sets the initial condition for movement onset, and neuronal activity evolve as a result of local cortical dynamics and inputs from other areas during movement execution [3], [4]. Rotational dynamics was observed after dimensionality reduction applied to MC population activity during arm reaching, revealing the basic patterns of complex single-neuron responses [5]. Recurrent neural networks (RNNs), which can approximate any dynamical system, is a widely used modeling tool to interpret how observed motor cortical patterns may emerge. Sussillo et al.[6] trained an RNN to generate muscle activity, using preparatory activity as initial conditions. Network with simple dynamics displayed surprising similarity to recorded MC data at both the single-neuron and population levels. From the view of pattern-generating network, Kao et al.[7] optimized the initial conditions for the desired movement, which bolsters the dynamical system perspective where initial conditions seed the subsequent movement. Moreover, external inputs are essential during movement execution. Kalidindi et al.[8] built an RNN model that receives sensory feedback from the arm, highlighting the sensory feedback in the motor cortical dynamics. However, most contemporary MC models are firing-rate-based networks that abstract away from individual spikes to firing rates[9], and neurons (often called nodes) in the models do not mimic the actual neural computations. Interpretation from simplified computational representations of neural activity may not capture the whole picture of motor cortical dynamics.

Alternatively, spiking neural networks (SNN) [10], [11], [12] are based on biologically plausible neurons and mimic the way neurons communicate through discrete spikes, enabling more accurate representations of temporal patterns and neural population dynamics. Dewolf et al.[13] presented an SNN model of the MC and cerebellum to control a three-link arm model for center-out reach task; their model can adapt to unknown perturbations and accounts for experimental data. Hennequin et al.[14] proposed a network with fine-tuned recurrent synapses and control different activity states for multidimensional movement patterns. The network displayed simultaneous firing rate and spiking variability. DePasquale et al.[9] proposed a training method for SNN by leveraging lower-dimensional factors extracted from either data or firing-rate-based networks. They trained the network to generate muscle activity recorded in the reach task, and spiking-neuron responses resembled to the real MC data. However, these models models that either rely entirely on external input or recurrent synapses or ignore sensory feedback during movement execution, could only partially capture the MC activity. Here, we propose a recurrent spiking neural network (RSNN) of MC, trained to control an arm model to perform reaching task as the monkey did (Figure 1A). Inspired by the multiple timescale model[15], the MC network is composed of two assembles of neurons, each with different temporal properties. The first assemble is composed of the “slow” neurons with large time constant, while the second is composed of “fast” neurons with small time constant. Multiple timescales can result in functional hierarchy in the motor control system, where basic elements are flexibly integrated into various patterns[15]. Moreover, the task input as well as proprioceptive feedback from the virtual arm went into the “slow” assemble, which interacts with “fast” assemble. Then the activity of “fast” assemble was readout into muscle activity that actuated arm model (Figure 1B, C). Despite not being fit to real MC data, our network reproduced neural population activity at both single-neuron and population-level, qualitatively and quantitatively. Our results suggest the possibility that multiple timescales is essential in motor cortical control.

## II. MODELING FRAMEWORK FOR REACH GENERATION

To simulate the process that monkey performs the reaching task, we elaborated a technique framework where an MC network controlled a musculoskeletal arm model (Figure 1). In this section, we describe the model architecture optimization. We first describe the MC network architecture, and introduce the single neuron and corresponding synaptic transmission model, and then illustrate how to transfer the neural activity to the hand movement. Besides, the definition of loss function and the training method are explained. Finally, we also described analyzing population dynamics of model and real MC data. All the simulations were performed on the BrainPy [16].

### A. The inputs to the network

The external inputs include the task input that represents target position, and sensory feedback. First, the target position in hand space was transformed into the desired joint angles (Figure 2). Second, we only used the proprioceptive feedback as the sensory feedback because during feedback control, human almost rely on the proprioceptive feedback to estimate hand motion[17], and multisensory integration display subadditive property[18]. The proprioceptive feedback ***u***_***p***_ consisted of joint angles and velocities. The input layer integrate all the input to

**Fig. 1.**
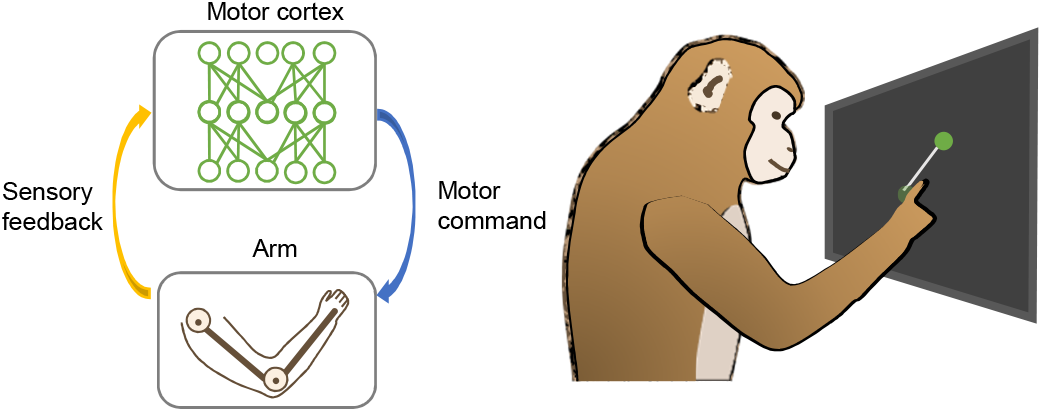
Understanding motor cortical control via neural network model. Monkey reaches his hand to the target on the screen. His brain receives instructions of the task, and generates motor command to the arm for movement. The sensory feedback was sent to the brain during reaching. The motor control system can be modeled as a closed-loop system. Motor cortex is simulated by artificial neural network, such as RNN and the arm is replaced by a musculoskeletal model.

**Fig. 2.**
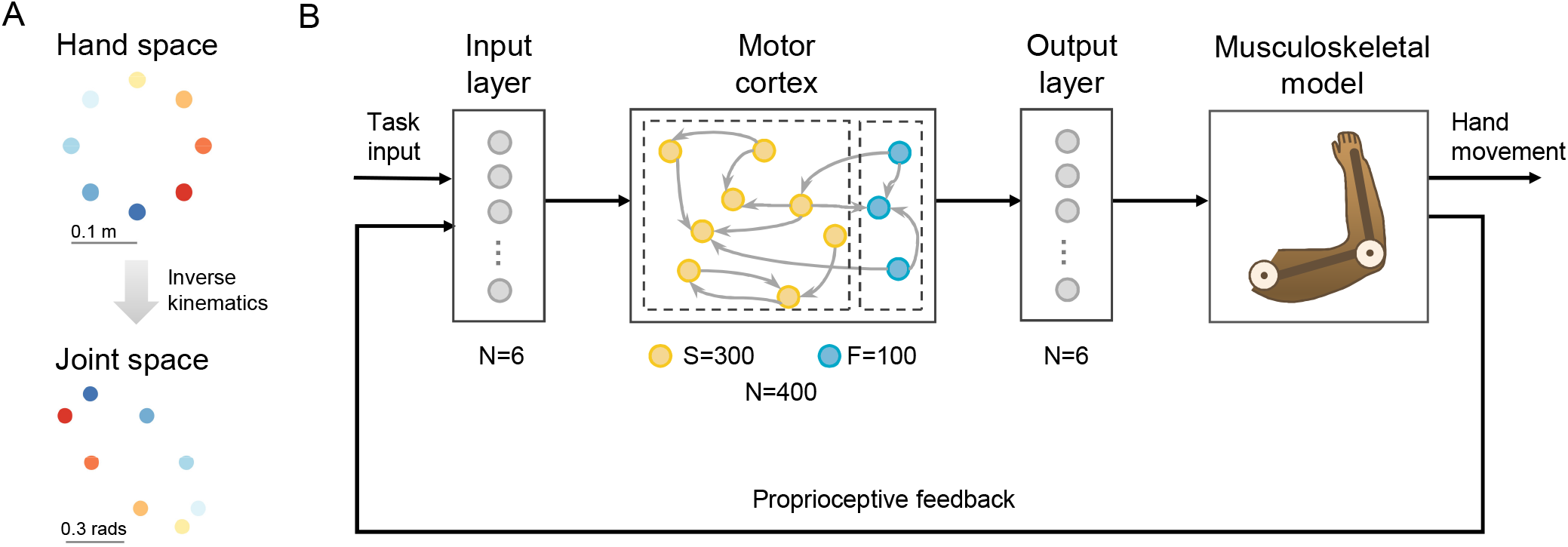
The structure of proposed Recurrent Spike Neural Networks. (A) Targets in cartesian and joint coordinates. (B) The task signal and feedback information are sent into input layer, a group of inter-connected spike neurons process and generate spike stream to driven the muscle to actuate the two link of arm.

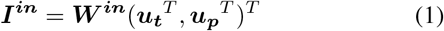

### B. Motor Cortex Network Model

#### 1) Single-Neuron Model

Single neuron was modeled as leaky integrate-and-fire (LIF) neuron[19], according to:

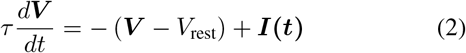

where *τ* = 8, and *τ* = 40 for “fast” and “slow” neuron, respectively. *V* is the membrane potential and *V*_rest_ = 0 is the resting membrane potential. When ***V*** reaches the threshold of 1 mV, the neuron emitted a spike. The membrane potential was then reset to -0.2 mV and holds for an absolute refractory period of 2 ms. ***I*(*t*)** is the time-variant synaptic inputs.

#### 2) Exponential Synapse Model

We assume that the release, diffusion, receptor binding of neurotransmitters and channel opening all occur instantaneously. The postsynaptic receptor conductance ***g*** was defined as:

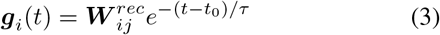

*τ* = 10 ms, *t*_0_ is the time of the pre-synaptic spike, *W*^*rec*^ reflected the conductance of neuron *j* to neuron *i*.

Most synaptic ion channels display an approximately linear current-voltage relationship when they open. Thus, the post-synaptic current is defined as:

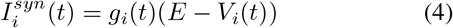

where *E* = 0 is the reversal potential, and *V* is the post-synaptic voltage.

#### 3) Spike Network Structure

Neurons (*N* = 400) in the MC network are divided into the fast (*N*_*f*_ = 100) and the slow (*N*_*s*_ = 300). The first assemble composed of spiking neurons with small time constant (*τ*_*f*_ = 5), resulting in rapid spiking, whereas the second assemble consisted of spiking neurons with large time constant (*τ*_*s*_ = 15), leading to slow spiking. All the neurons have recurrent connections to each other at both within assemble and assemble to assemble. Only half of the presynaptic neurons projected to postsynaptic neurons for sparse connection. The “slow” assemble receives input and “fast” neurons’ spikes are mapped into muscle activity:

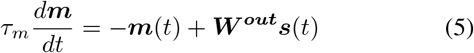

where *τ*_*m*_ = 200 for smooth, ***m*(*t*)** is the six-dimensional muscle activity, ***s*** is the spikes from the slow assemble, ***W*** ^***out***^ is the output matrix.

### C. Musculoskeletal Arm Model

To simulate arm reaching, we constructed a planar two-link arm model as detailed in the previous work [20], [21]. Briefly, the upper link and the lower link are connected at the elbow (Figure 1). The arm model is actuated by six muscles, whose activity is converted to muscle forces according to the length/velocity properties of muscle. The muscle forces are further converted to the joint torques ***τ* (*t*)** that changes the arm state:

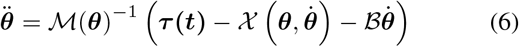

where the arm state *θ* includes the joint angles and velocities 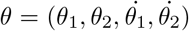 . The non-linear function *ℳ* and *𝒳* is the matrix of inertia and the centripetal and Coriolis forces. The position of the end effector (hand) is:

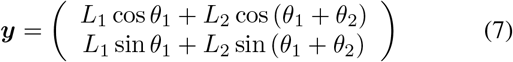

### D. Network Optimization

Network was optimized to control the arm model to reach the target after go cue within in 700 ms. The loss function is the squared error between the actual hand position and target:

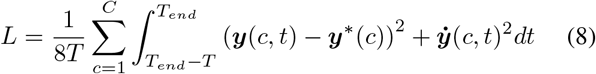

where ***y***^*∗*^ is the target position, equally spaced in eight directions on a circle with a radius of 10 cm, 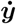 is the hand velocity. *T*_*end*_ = 1200 and *T* = 500. The network weights were trained by back-propagation. We used Adam optimizer with learning rate of 10^*™*4^ and iterated it 2000 times.

### E. Statistical Analysis

#### 1) Neuronal data analysis

We analyzed the recordings from a monkey’s MC during reaching [5] and spikes data from our model. As for the monkey data, we extracted the activity of 202 neurons in eight straight reach conditions. We processed the spike trains by computing the firing rates and averaging them for each reach condition. Then we smoothed firing rates by a 60 ms Gaussian filter. All the firing rates were computed for the execution period, time-locked to the movement onset. The following analyses were based on firing rates.

#### 2) jPCA

jPCA captures the rotational dynamics within firing rate responses[5]. First, data was reduced to ***X***_***red***_ *∈* ***R***^k*×ct*^ by principal component analysis (PCA), where c = 8 is the number of reach conditions, and t = 700 is the number of time points, k = 6 is the number of principal components. Second, the reduced data was fitted into a linear dynamical system ***X***_red_ = ***M*** _*skew*_***X***_red_, where ***M*** _skew_ is a skew-symmetric matrix that has purely imaginary eigenvalues. The neural trajectory in the plane of the first two eigenvectors of the ***M*** _*skew*_ rotates most strongly.

## III. EXPERIMENT AND DISCUSSION

### A. The comparison of model and data

The optimized model can simulate the MC and had a good performance in the eight-direction center-out reaching task(Fig. 3). Both fast and slow neurons in the RSNN displayed strong directional tuning in the raster plot and PSTH and temporal complexity(Fig. 4 A and B): the slow example neuron only spiked at the movement onset and the fast example neuron fired during the whole movement epoch with multiphase and heterogeneous firing pattern which was also observed in monkey’s MC. At neural population level, we computed the low-dimensional patterns via jPCA. All the neural population responses displayed rotational dynamics(Fig. 4 C). Although the fast assemble with fewer neurons than the slow assemble, its rotational structure was more similar to the overall dynamics, suggesting that the neurons connected to the output layer contribute more to the neural population responses. Furthermore, the ratio of fitting to the constrained and unconstrained dynamical system was 0.80, which indicates that the majority of the output layer’s dynamics captured more rotational dynamics. To quantify the similarity between real and model patterns, we first reduced the dimensionality of the two datasets by PCA, obtaining the top 18 (monkey), and 20 (model) principal components, and then performed canonical correlation analysis (CCA) in a 600 ms window starting after movement onset. The average correlation is 0.89.

**Fig. 3.**
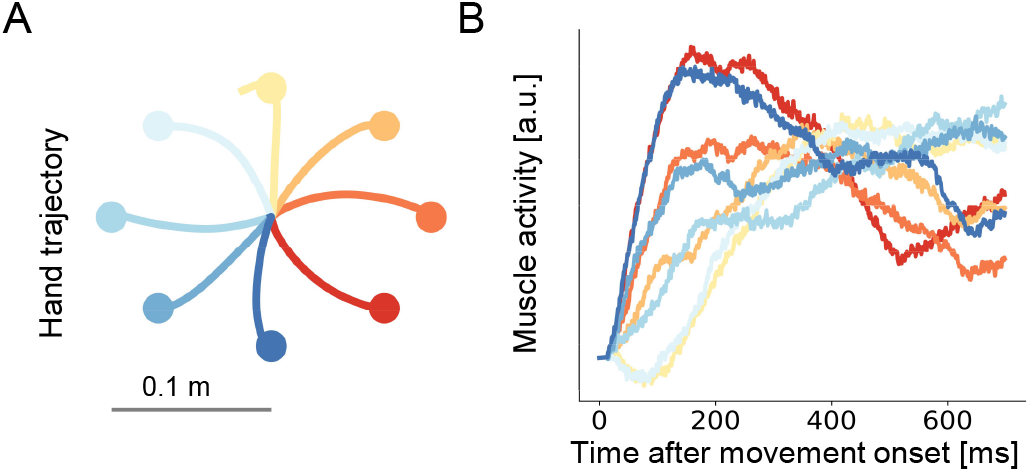
Hand trajectory (A) and muscle activity (B) produced by the model.

**Fig. 4.**
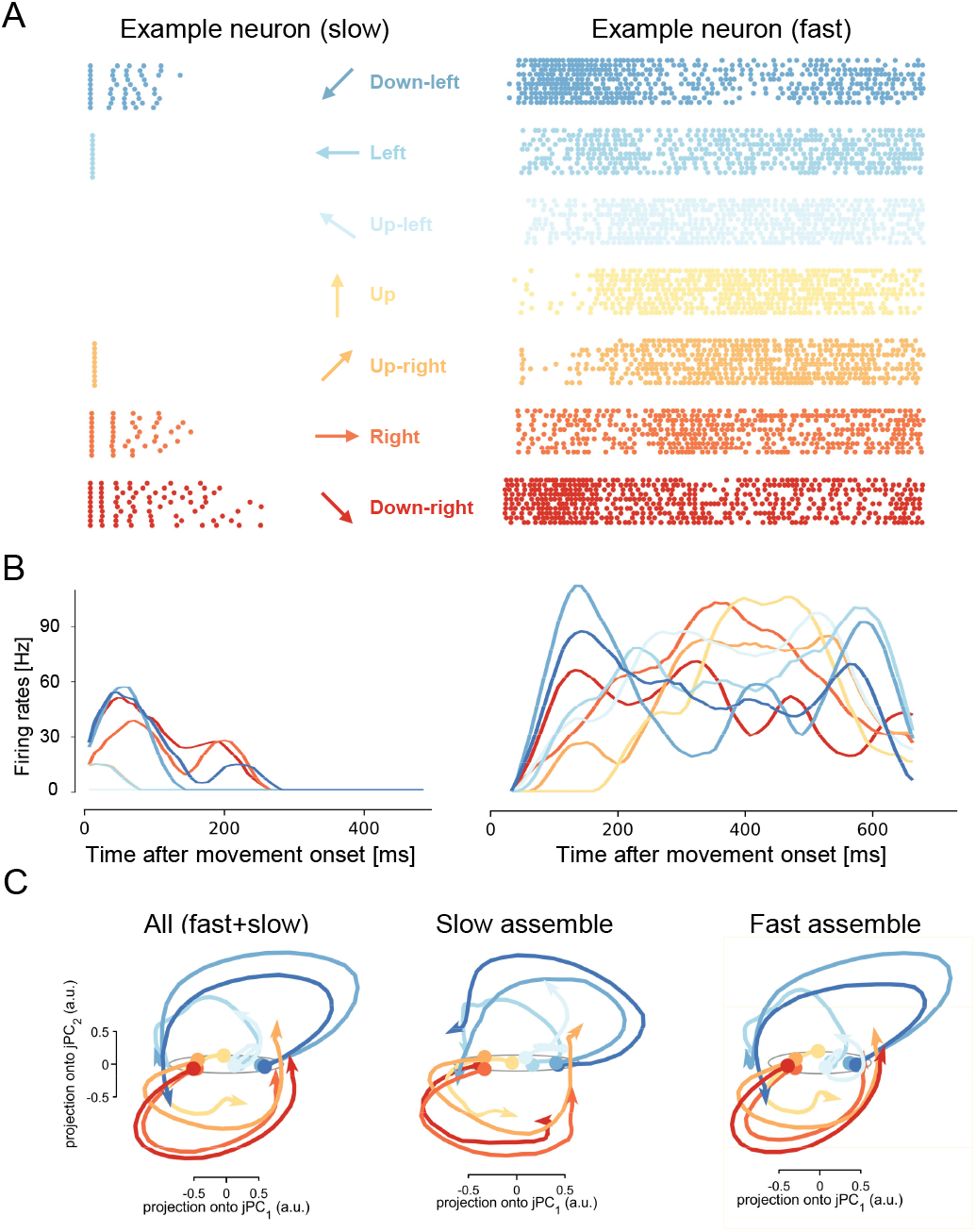
Neural activity in the model. (A) Raster plots of spikes from two example neurons for 11 trials per direction. (B) Trial-averaged firing rates of the example neurons. (C) Neural population dynamics of all, “fast” and “slow” neurons after jPCA.

## IV. CONCLUSION

In this paper, we proposed an recurrent spike neural network based modeling approach of motor cortex, which controls an arm model to perform the center-out reaching task. Experiments show that our model reflects the real motor cortical activity at both both the single-neuron and population levels, qualitatively and quantitatively, suggesting the possibility that motor control is driven by neurons with different time constants in the MC.

In the future research, we will optimize the network structure and training efficiency, increase the number of neurons, explore the mechanism of spike accumulations, and apply the recurrent spike neural network to perform a wide variety of tasks.

## ACKNOWLEDGMENT

This work was supported by the STI 2030—Major Projects (2022ZD0208604), and the NSFC (62176151 and 61773259). We are grateful to Prof. Xinyu Chai and Prof. Liming Li for instruction, to BrainPy team for computation support, to Dr. Shenjiang Rong and Dr. Jieji Ren for discussion.

